# Three-Dimensional Mechanical Cooperativity Optimises Epithelial Wound Healing

**DOI:** 10.1101/2025.07.18.665363

**Authors:** Shu En Lim, Pablo Vicente-Munuera, Robert J. Tetley, Martin Zhang, José J. Muñoz, Yanlan Mao

## Abstract

Epithelial tissues serve as critical physical barriers that protect organs, making efficient repair essential upon damage. To achieve this, tissues must quickly react, forcing cells to rearrange and accommodate. Therefore, cells must cope with forces that will deform their shape to achieve these sudden but necessary changes. In the *Drosophila* wing disc, we showed how tissue fluidisation accelerated wound healing in 2D. However, the 3D aspect of tissue repair remains poorly understood. Here, we uncover a new mechanism aiding the tissue in repairing itself by changing cell height. We find actomyosin contractile cables at the wound edge connecting the apical and basal cell surface, which indent the apical side of the tissue and deform the basement membrane (BM), respectively. To understand the role of the different repair mechanisms, we developed a 3D vertex model allowing apico-basal intercalations. The model predicts that lateral cables play a role in regulating cell-cell intercalations, confirmed by *Drosophila* mutations affecting cell deformations. Our results demonstrate that lateral cables cooperate with the apical purse string to drive 3D cell shape changes and intercalations to promote more efficient wound repair.

## Introduction

Effective wound healing relies on robust, tightly coordinated responses that depend on the mechanical properties of cells and their surrounding environment [1,2]. Most studies relating to tissue mechanics and wound healing have considered monolayered epithelial tissues as two-dimensional (2D) planar structures [3]. The three-dimensional (3D) shape of epithelial cells, particularly the organisation and mechanics of their lateral domains, plays a crucial role in driving tissue morphogenesis. For instance, developmental tissue folding in *Drosophila* wing and leg imaginal discs has been shown to rely on apicobasally orientated forces driven by lateral accumulations of myosin II [4,5]. While there is some evidence that epithelial cells change shape in 3D during wound healing, mainly in flat cell culture epithelia, how these 3D shape changes are controlled and whether they contribute to epithelial wound healing has not been explored [6–9].

In addition to intrinsic cell mechanics, 3D tissue and cell shape are also driven by communication to the external environment via cell-BM adhesions, for example, in the *Drosophila* follicular epithelium [10]. Cells also transmit forces to the underlying substrate via focal adhesions during wound closure, as quantified using traction force microscopy of MDCK cells [1]. Furthermore, Integrin-mediated adhesion is remodelled during wound closure in *Drosophila* embryos [11]. Despite this, the relationship between 3D cell shape and substrate adhesion during wound healing has not yet been investigated.

In parallel, 2D vertex models have been widely used to study tissue mechanics during wound healing and other biological processes, helping to understand how forces at cell interfaces shape tissue behaviour [2,12–14]. Such models have shown that, depending on the initial cell geometry and tissue mechanical properties, distinct closure mechanisms can arise to seal the wound [2,15,16]. However, extending these models to 3D has proven challenging [8,17], particularly when accounting for complex spatial features such as apico-basal intercalations or scutoids [18]. A previous 3D hybrid model incorporating differential contractility at apical, basal, and lateral surfaces demonstrated that an apical purse-string can drive wound closure when combined with volume preservation [8].

To explore these questions in a physiologically relevant context, the *Drosophila* wing disc offers an ideal *ex vivo* system for investigating 3D cell behaviours. The wing disc comprises approximately 45μm tall pseudostratified columnar epithelial cells encased by a basement membrane (BM). We have previously characterised an actomyosin purse string similar to that found in wounded *Drosophila* embryos [19] and shown how tissue fluidity plays key roles in enabling intercalation for scarless wound closure in the wing disc [2]. Here, we use live imaging and laser ablation of the wing disc to characterise 3D cell shape changes during wound healing. We have discovered a novel actomyosin-driven lateral cable mechanism independent from the apical purse string. By combining genetic perturbations with an intercalation-enabled 3D vertex model, we reveal how the distribution of myosin-II-driven forces dependent on substate adhesion can influence 3D mechanisms of wound repair.

## Results

### Wing Disc Wound Healing Is Accompanied By 3D Cell And Tissue Shape Changes

We used wing discs between 96-120 hours after egg laying because, at this stage, the pouch region is relatively flat and thus easier to wound apically (Fig. 1A). Following wounding by laser ablation and live time-lapse imaging, we observed that the wound extended the entire height of the epithelium, from the apical surface to the basal surface (Fig. 1B). This led us to examine cell and tissue behaviours along the tissue’s apicobasal axis during wound healing. 120 min after wounding, we consistently observed that the apical and basal surfaces had indented towards the centre of the wound, forming distinctive funnel-like shapes (Fig. 1C). We found that the tissue height at the wound edge decreases by approximately 40% over 120 min of closure (Fig. 1D). These observations indicated that wound edge cells were shortening as healing progressed leading us to hypothesise that such 3D cell shape changes could facilitate wound closure. Specifically, as cells shorten, they expand their planar area into the wound, reducing the wound gap. This hypothesis relies on wound edge cells conserving their volume during shortening. Previous studies of *Drosophila* embryonic epidermis wounds have reported differing results, with some showing that wound-edge cells conserve volume [7] whilst another reports an increase in volume [20]. To quantify cell volume, we induced sparse 1-2 cell cytoplasmic mCherry-expressing clones in the wing disc and wounded tissue so that these clones were on the wound edge (Fig. 1E, S1A). We quantified the clone volume and height over the first 120 min of closure and found that while the clones decreased in height, they did not significantly change in volume (Fig. 1F-G, Supplementary Fig. 1B), supporting our hypothesis that cell shortening at the wound edge could facilitate wound closure.

**FIGURE 1:**
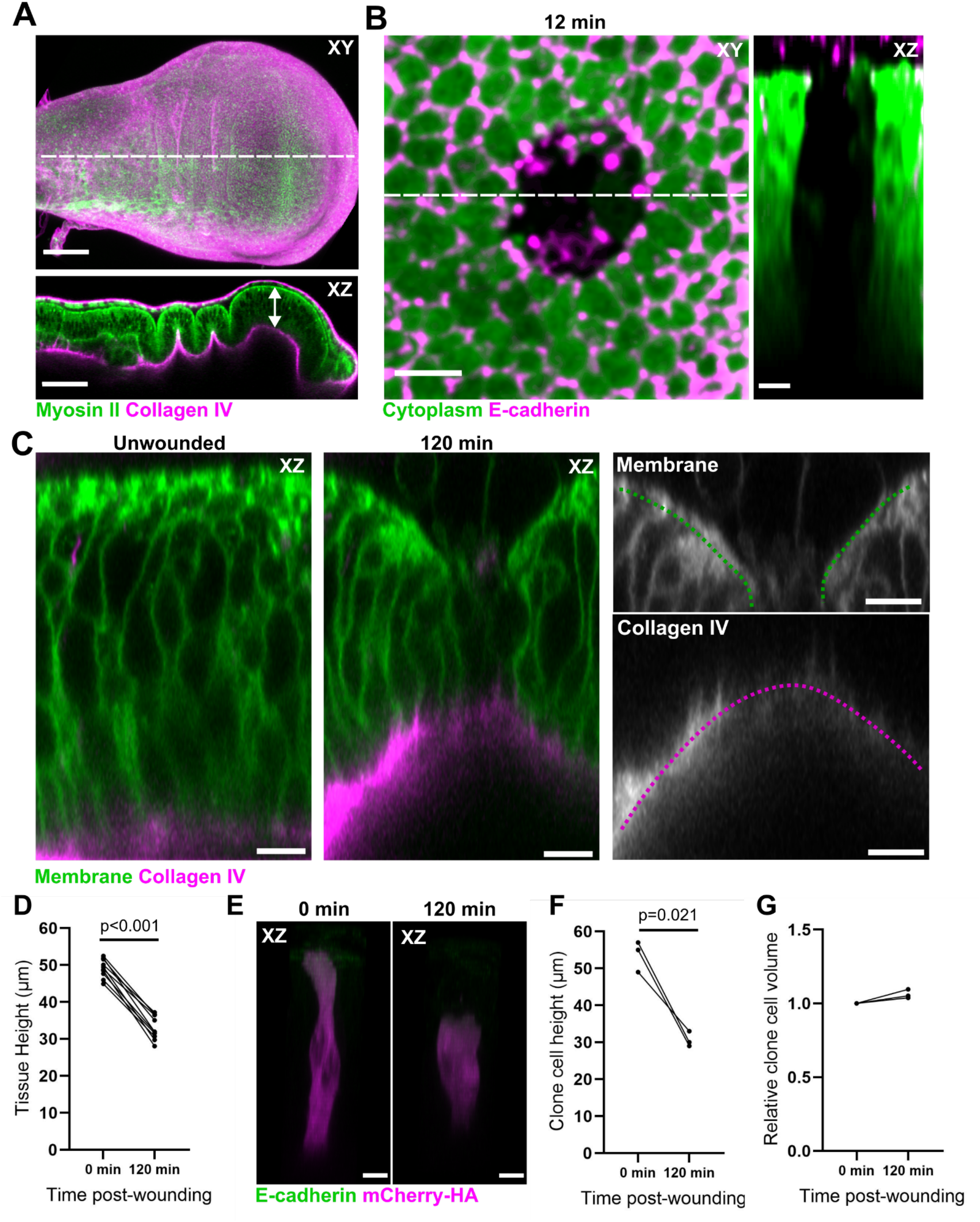
3D cell and tissue shape changes characterise wing disc wound healing. A) Third instar Drosophila wing discs are composed of columnar epithelial cells surrounded by a thin layer of BM (collagen IV, magenta). These cells are enriched apically by myosin (green). Dotted line indicates the position of the XZ view below, with an arrow marking the 45μm height of the wing disc. Scale bars, 20μm. B) Wounds in the wing disc propagate through the whole tissue 12 min after wounding. Dotted line indicates the position. of the adjacent XZ view. Scale bars, 5µm. C) Wing disc expressing GFP-CAAX (green) and vkg-mScarlet (magenta) showing a cross section of an unwounded region compared to the wound edge 120 min after wounding. Scale bars, 5μm. D) Tissue height before wounding and 120 min after wounding (n=10). E) Cytoplasmic mCherry 2-cell clone with adherens junctions marked with GFP-tagged E-cadherin. Scale bars, 5μm. F) Clone cell height (n=3). G) Clone cell volume normalised to initial volume (n=3). For D, F and G, plots show individual data points with the corresponding paired values connected by a line and p-values from ratio paired t-tests.

### Contractile Lateral Actomyosin Cables Form at the Wound Edge and Deform the Tissue

To investigate how wound edge cells change shape in 3D, we used 3D timelapse imaging to quantify wound edge dynamics. We marked the wound edge using GFP-tagged myosin II, the primary force generator in this tissue, which is enriched at the apical surface (Fig 2A) [2]. Imaging in 3D, we found that wound edge cells also begin indenting within 5 min and the indentation depth increases linearly over time (Fig. 2B).

**FIGURE 2:**
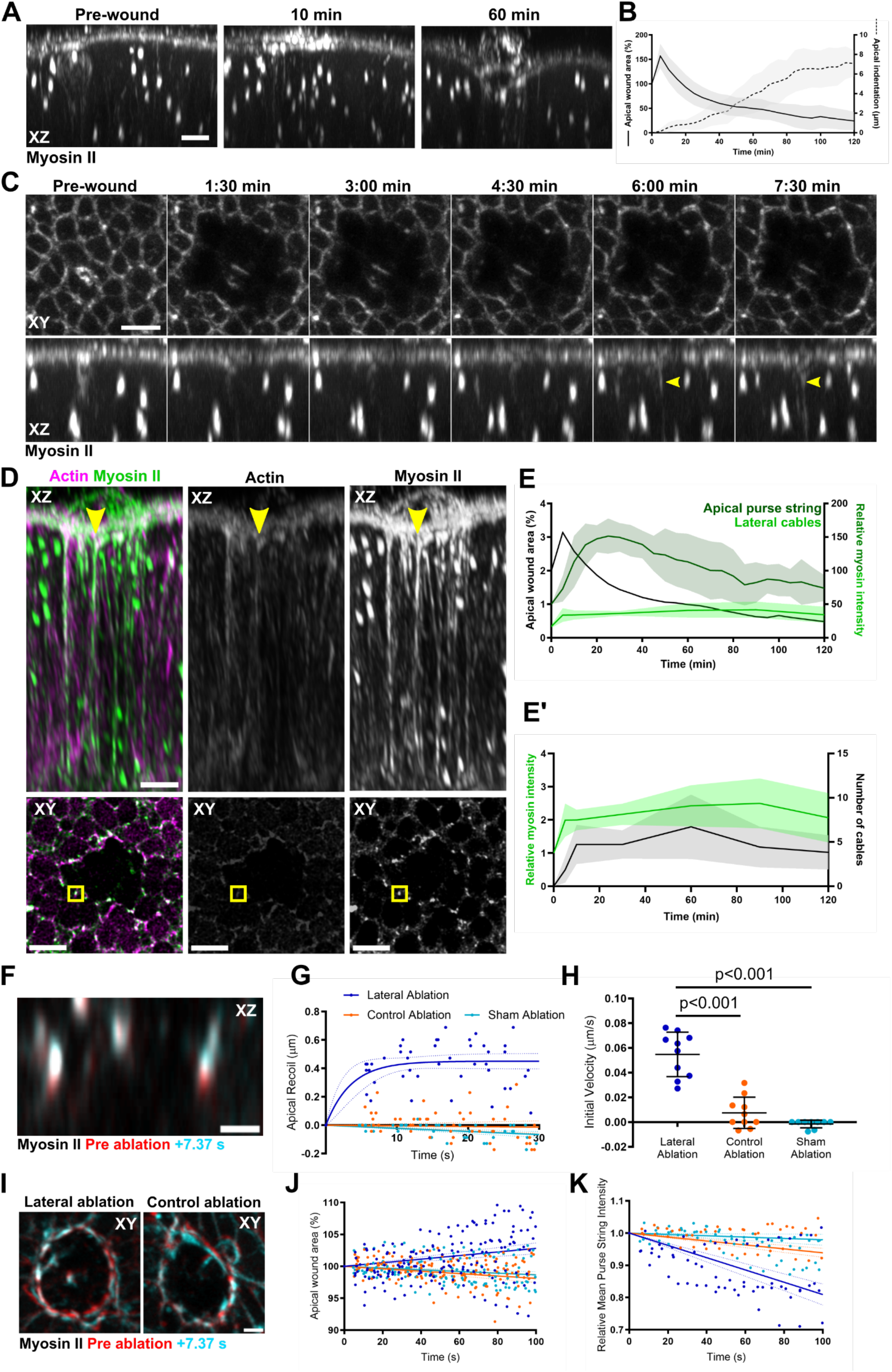
Lateral actomyosin cables form at the wound edge and deform the tissue via increased apicobasal contractility. A) Timelapse images of wound cross sections with myosin II-GFP in grey. Scale bar, 5µm. B) Quantification of apical wound area (solid line) and apical indentation (dotted line) over the first 120 min of closure. C) XY and XZ max projections of a timelapse movie of a wounded wing disc expressing sqh-mKate to label myosin II (n=7). D) Cross-sectional max projection and XY plane of a wound in a wing disc expressing sqh-GFP in green and LifeAct-mCherry in magenta 60 min post-wounding. The yellow arrow and box indicate a lateral cable. Scale bars, 5µm. E) Apical wound area (black), relative apical purse string myosin II intensity (dark green) and relative lateral cable myosin intensity (light green) over the first 120 min of wound closure (n=7, from B). E’) Relative myosin intensity of the lateral cables normalised to the baseline lateral myosin II intensity before wounding. F) Cross-sectional view of the apical purse string marked with myosin II-GFP (as marker of apical surface) pre-lateral ablation (red) and 7.37s post-lateral ablation (light blue). Scale bar, 5µm. G) Apical recoil of the purse string myosin II from lateral ablation (dark blue, fitted with one phase decay), control ablation (orange, fitted with a straight line) and sham ablation (light blue, fitted with a straight line). Dotted lines indicate 95% confidence intervals. H) Initial velocity. I) Apical purse string, pre-ablation (red), post ablation (blue). J) Apical wound area relative changes after lateral cable ablation. Scale bar, 5µm. K) Relative Mean Apical Purse String intensity changes after lateral cable ablation. For G, H, J and K; n=10.

The sustained apicobasal shortening at the wound edge suggests the presence of an active force-generating mechanism initiated rapidly upon wounding. At the apical surface of the wound edge, we observed strong myosin II accumulation forming a classic purse string, consistent with previous reports [2,19,21]. When we examined deeper z planes basally through the wound, we noticed myosin II accumulations decorating the lateral edges of wound edge cells, emerging approximately 6 min after wounding (Fig. 2C). On closer inspection, we could identify a single cable in most cells, which was closely localised to sites of former tricellular junctions and formed a continuous structure with the apical purse string (Figure S2A-B). Using high-power, single-timepoint imaging of live tissues, we could resolve the full extent of these cables, revealing they spanned the entire apicobasal axis and colocalised with actin (Fig. 2D). Based on their localisation, orientation, and composition, we refer to these structures as lateral cables.

Due to phototoxicity, it was not feasible to continuously image GFP-tagged myosin II along the full length of the lateral cables at high laser power and with high spatial-temporal resolution. We therefore focused on the apical portion of the lateral cables where signal intensity was strongest, and quantified myosin II intensity throughout wound healing. Compared to baseline apical myosin II intensity, lateral cables exhibit significantly lower intensity (Fig. 2E). However, compared to baseline lateral myosin II intensity before wounding, lateral cable myosin II intensity increases by 50% within 5 min post-wounding and then remains stable (Fig. 2E’). In contrast, the apical purse string sharply increases, reaching nearly 3-fold the baseline apical myosin intensity before gradually declining as healing progresses (Fig. 2E).

Given their geometry and continuity with the apical purse string, lateral cables were strong candidates for generating vertical tension at the wound edge to shorten cells. To test their role in force generation, we performed targeted nano-ablations of the cables and quantified the impact on apical indentation (Fig. 2F). Simultaneous severing of all lateral cables (lateral ablation, Fig. S2C) caused the apical surface to recoil upwards, whereas nano-ablation of cortical regions away from the lateral cables (control ablation, Fig. S2C) caused minimal recoil (Fig. 2G). We discarded nano-ablation experiments where cells were killed (Supplementary Methods). In the absence of any ablation (sham ablation), we observed a small increase in apical indentation depth, as would be predicted if proceeding closure was unaffected (Fig. 2F-H). Overall, specific severing of lateral cables produced significantly higher recoil velocities (Fig. 2H). These results suggest that the apicobasal axis is placed under increased tension by the lateral cables, which indent the apical surface.

Following nano-ablation, we also quantified both the apical wound area and purse string myosin II intensity (Fig. 2I-K). Lateral ablation led to an increase in apical wound area (Fig. 2J) and a decrease in mean myosin II intensity in the apical purse string, whereas neither effect was seen in control and sham ablation (Fig. 2K). Together, these findings suggest that lateral actomyosin cables contribute to the mechanical stability and maintenance of wound closure speed as well as the apical purse string during wound healing.

### A 3D Vertex Model Predicts Cooperative Forces Accelerate Wound Closure

To decouple the individual contributions of the apical purse string, lateral cables and intercalation during wound healing in the *Drosophila* wing disc, we developed a novel 3D vertex model. To create the initial geometry of the tissue, we started from a segmented flat 2D image of the wing disc epithelium [22] which we extruded to generate a full 3D tissue geometry with a height equivalent to 45μm. A key innovation of our model is the ability to support intercalations in 3D, allowing for dynamic changes in apico-basal topology and the emergence of scutoids (Fig. 3A, and Supplementary Methods). When a T1 transition occurs on either the apical or basal surface, a scutoid often forms. To account for this, the model dynamically generates a new face between the two cells involved, localised only to the surface domain where the T1 takes place (Fig. 3A, green arrow).

**FIGURE 3:**
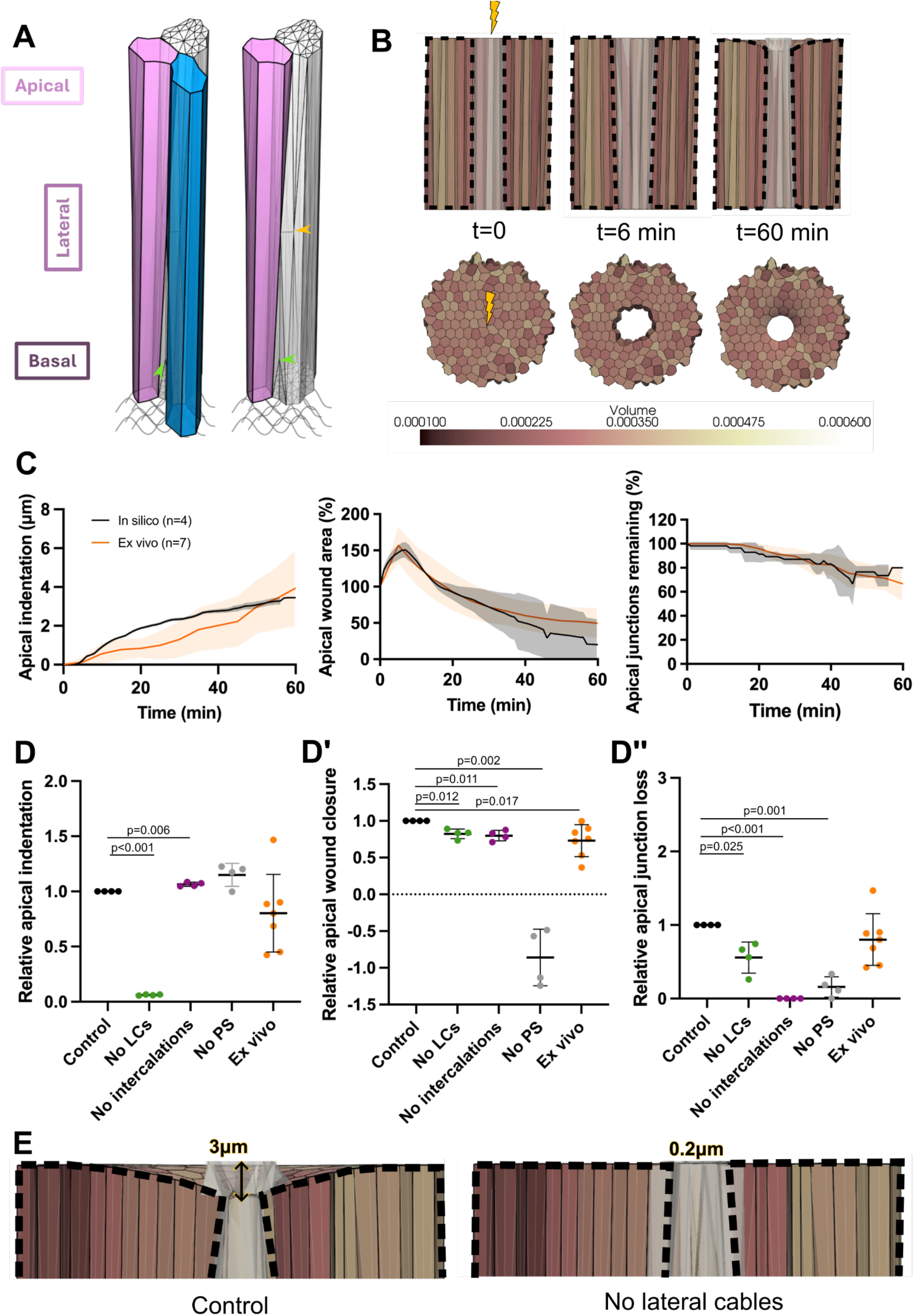
Computational model predicts different wound dynamics in the absence of purse string, lateral cables or intercalations. A) Representation of cells in the model. Left shows three cells forming an apico-basal intercalation. Right shows two cells with the scutoid face more visible. Yellow arrow is the face centre that connects two cells without an apico-basal transition between them. Green arrow is a face centre representing a scutoid face connecting two cells only at the apical side. Note that white cell displays the discretization in the model, showing all the vertices and their edges in black. Grey curved lines represent the BM. B) Representative tissue dynamics over time after ablation at t=0. At t=6, the tissue recoils only apically, increasing its wound area. At t=60, the tissue closes the wound and indents. Debris cells are represented with pale white colours. Cell colour scale indicates their volume. Black dotted lines highlight the tissue contours. C) Wound healing dynamics of in silico simulations (black, n=4) and ex vivo movies (orange, n=7) for 60 min. Apical wound area percentage after wounding; apical indentation; and percentage of apical junctions remaining. Shaded areas represent the standard deviations (SD). D-D’’) Effect of the absence of a mechanism on the velocities of the apical indentation (D), apical wound area (D’), and apical junctions loss (D’’) normalised with its control simulation (black) at 60 min post-wounding. Ex vivo velocities are normalised using the average of the in silico controls. ‘No LCs’ stands for ‘no lateral cables’ (green), and ‘No PS’ means ‘no purse string’ (grey). n=4 for all simulations, and n=7 for ex vivo. Individual data points are shown with the mean (horizontal line), SD (bars) and p-values from two-tailed Mann-Whitney tests. E) Zoom in of apical indentation on control vs ‘No LCs’ simulation at end time. The arrow corresponds to the average measured apical indentation of the n=4 simulations.

Since, initially in our model, both apical and basal surfaces have the same topology, we performed several intercalations in the basal surface to introduce topological differences between the apical and basal surfaces. This generated an initial state where approximately 50% of cells were scutoids, with a more regular polygonal distribution apically than basally (Fig. S3A). Each cell is modelled as a 3D polyhedron with bulk and surface area elasticity, a substrate attachment term and a cell shape deformation term (Fig. 3A, and Supplementary Methods). We included additional line tension terms to model the purse string and lateral cables as independent actomyosin pools. In the resting state, all line tensions are set to zero. Because the geometry derived from segmentation may not be mechanically equilibrated, we first simulated vertex dynamics until the tissue reached a steady-state pre-wound configuration (Fig. 3B, t=0, Supplementary Methods).

We then performed an *in silico* wounding experiment by ablating 10 cells in the middle of the tissue (Fig. 3B), modelling them as debris with altered mechanical properties (Supplementary Methods). The model was parameterised using experimental data to reproduce key features of the experimental wound healing data observed *ex vivo*. These included strong recoil at approximately 6 minutes after the ablation, and apical indentation, intercalation and apical wound area closure over the first 60 minutes (Fig. 3C). Further, we used experimentally measured myosin II intensities of purse string and lateral cables as a proxy for their contractile strength (Fig. 2E) [23]. We could replicate common 2D mechanical features (n=4 simulations with different topologies) such as peak recoiling at 6 minutes, apical wound area percentage decay rate, and apical junction loss rates as those observed in the *ex vivo Drosophila* wing disc during the first 60 minutes post-wounding (Fig. 3B-C, and Supplementary Movie 1).

Given the experimental complexity of selectively perturbing individual contributions *ex vivo*, particularly distinguishing between spatially distinct myosin II pools, we used our novel computational model to predict wound healing dynamics in the absence of the apical purse string, lateral cables and intercalations, respectively (Fig. 3D-D’’). Using four different control geometries, we ran simulations for each perturbation and compared wound healing by normalising closure velocities. All the velocities were normalised with regard to their corresponding control geometry simulation.

The strongest impairment in wound healing occurred when the purse string was removed, indicating that it is the primary driver of tissue repair (Fig. 3D’, and Supplementary Movie 2). This aligns with its role as a contractile cable acting predominantly in the X–Y plane [1], whereas lateral cables generate force along the apicobasal axis (Fig. 3E). Notably, eliminating either lateral cables (Supplementary Movie 3) or intercalations (Supplementary Movie 4) also significantly reduced healing efficiency, consistent with model predictions showing a complete loss of apical indentation in the absence of lateral cables (Fig. 3D and Fig. 3E). Unexpectedly, the model also revealed that removing lateral cables impaired cell intercalation (Fig. 3D’’), suggesting that these structures may facilitate tissue remodelling by reducing cell height while preserving cell volume. This highlights a previously unappreciated role for lateral cables in coordinating 3D tissue architecture during wound closure.

### Lateral Cable Assembly and Apical Indentation Occurs Independently of the Apical Purse String

To experimentally test whether lateral cable formation is independent of apical purse string myosin II recruitment, we used a temperature-sensitive *shibire* (*shi*^*TS*^) mutant to block dynamin-mediated endocytosis (Fig. 4A), which has previously been shown to prevent purse string assembly in *Drosophila* embryo wounds [24]. As expected, wound closure slowed 2.5-fold (Fig. 4B-B’) and apical purse string myosin II intensity was reduced by approximately 40% during the first 60 min of closure in *shi*^*TS*^ wing discs, (Fig. 4C-C’). Loss of apical junctions at the wound edge, which reflects cell intercalation events, was unaffected during this early stage of closure (Fig. 4D-D’) [2]. Interestingly, apical indentation rate (Fig. 4E-E’) and initial lateral cable formation (Fig. 4F-G) were unaffected, indicating that lateral cable formation and cell shortening can occur independently of the apical purse string. Fewer lateral cables remained after 60 min (Fig. 4F), while overall myosin II recruitment to individual lateral cables was not significantly altered (Fig. 4G). This indicates that myosin II stabilisation in lateral cables may require cooperation with the apical purse string.

**FIGURE 4:**
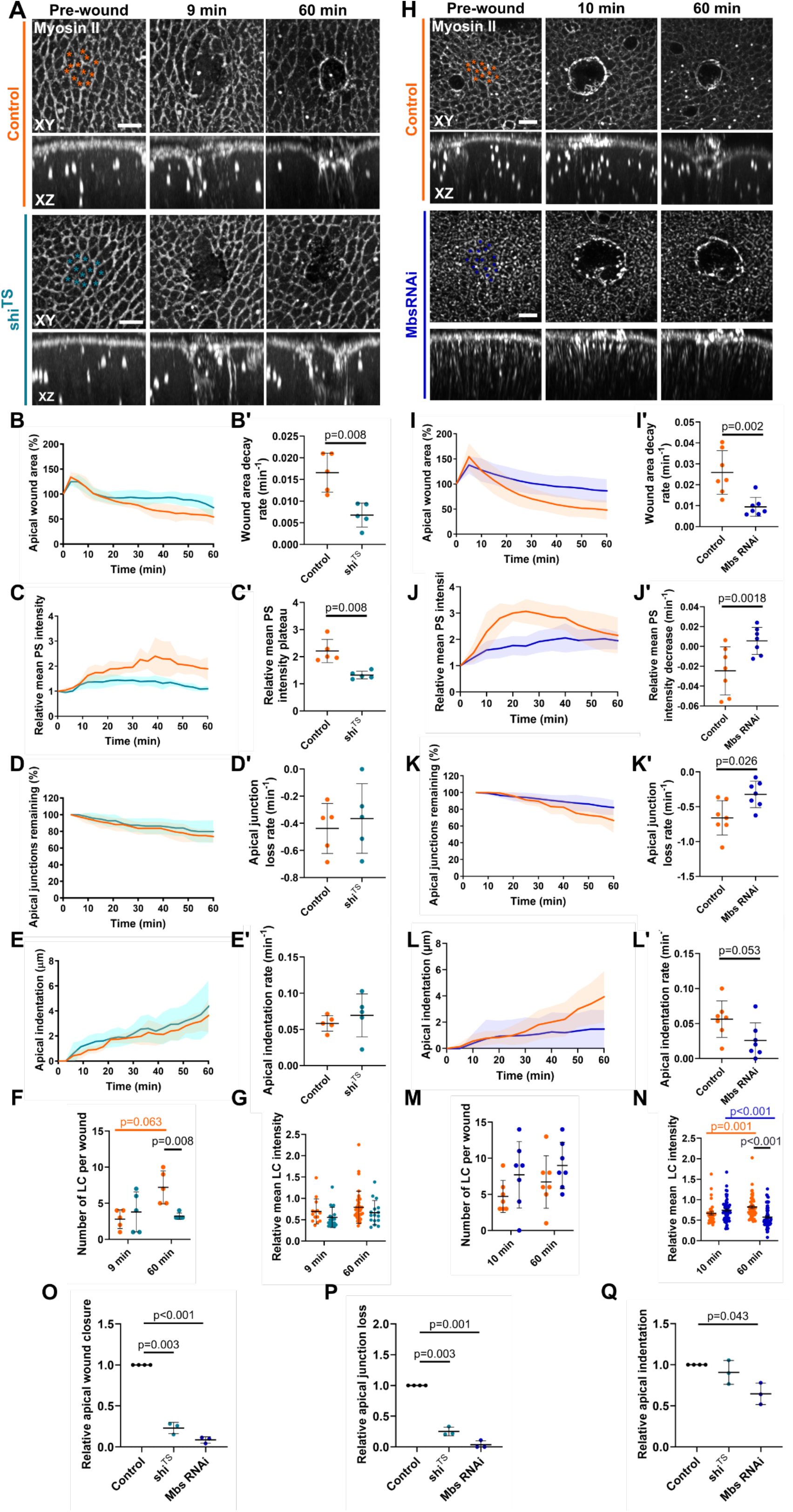
Regulated myosin activity is required for effective 3D shape change. A) Timelapse images for temperature shifted wounded control discs (orange, n=5) and wounded shi^TS^ discs (blue, n=5) expressing sqh-GFP. Asterisks indicate dead cells. Scale bar, 5μm. B) Apical wound area, 100% = original wound area. B’) Wound area decay rates from fitting one-phase decay curves of individual replicates from B. C) Relative PS myosin II intensity. C’) Relative PS myosin II intensity plateaus from fitting one phase associations of individual replicates from C. D) Wound edge apical junctions remaining as cells are eliminated from the wound edge. D’) Junction loss rates from fitting straight lines of individual replicates from D. E) Apical indentation. E’) Apical indentation rates from fitting straight lines of individual replicates from E. F) Number of cables per wound 10 min and 60 min after wounding. G) Relative mean LC myosin II intensity 10 min and 60 min after wounding. H) Timelapse images for wounded control discs (orange, n=7) and wounded Mbs RNAi discs with genotype (blue, n=7) expressing sqh-GFP. Asterisks indicate dead cells. Scale bar, 5μm. I) Apical wound area. I’) Wound area decay rates from fitting one-phase decay curves of individual replicates from I. J) Relative PS myosin II intensity. J’) Relative PS myosin II intensity reduction rates from fitting straight lines of individual replicates from 20 min post-wounding from J. K) Apical indentation. K’) Apical indentation rates from fitting straight lines of individual replicates from K. L) Apical junctions remaining as cells are eliminated from the wound edge. L’) Junction loss rates from fitting straight lines of individual replicates from L. M) Number of cables per wound 10 min and 60 min after wounding. N) Relative mean LC myosin II intensity 10 min and 60 min after wounding. O) Relative apical wound area velocity, P) relative apical junction loss, and Q) relative apical indentation velocity regarding control at 60 minutes. n=4 for control, n=3 shi^TS^ and Mbs RNAi in silico mutations (Supplementary Methods). For B-E and I-L, shaded areas represent the SD. For B’-E’, F-G I’-L’ and M-N, plots show individual data points with the mean (horizontal line), SD (bars) and p-values from two-tailed Mann-Whitney tests. For F, p-values between different timepoints were calculated using Wilcoxon matched-pairs signed rank tests. Scale bars are 5 µm.

To explore an alternative method to manipulate the tension of the purse string, we next asked how wound healing is affected when myosin II activity is hyperactivated. Previous work has shown that myosin II hyperactivation by Mbs RNAi balances the tissue tension between resting unwounded cells and the purse string [2]. We find that hyperactivation of Myosin II via Mbs RNAi significantly impaired healing dynamics compared to controls (Fig. 4H). By 60 min, Mbs RNAi wounds remained 100% open, whereas control wounds had closed to 50% of their initial area (Fig. 4I). The wound closure rate in Mbs RNAi wounds was approximately 2.7-fold slower (Fig. 4I’).

The dynamics of the apical purse string were also significantly altered, with apical purse string myosin II intensity peaking 2-fold above resting levels in Mbs RNAi wounds, compared to 3-fold in controls (Fig. 4J). While myosin II intensity then gradually declined in control wounds, it remained constant in Mbs RNAi wounds, indicating prolonged myosin II stability (Fig. 4J’). This reduced response likely reflects constitutively elevated myosin II levels due to impaired myosin phosphatase activity, consistent with our previous findings that Mbs RNAi elevated both purse string and resting tensions to maximum levels [2].

In line with our earlier work, wound edge apical junction loss was significantly decreased by more than half (Fig. 4K-K’). Additionally, 3D cell shape changes were impaired, with apical indentation showing a decreasing trend and occurring approximately 3.2-fold more slowly in Mbs RNAi wounds (Fig. 4L–L’). Although the number of lateral cables per wound did not differ significantly (Fig. 4M), their mean myosin II intensity declined by approximately 20% after 60 min in Mbs RNAi wounds (Fig. 4M-N). This intensity was also significantly lower than controls at 60 min (Fig. 4N).

Although lateral cables still formed, the declining myosin II intensity over time suggests that dynamic turnover is required to maintain their stability and contractility to drive apical indentation. By hyperactivating myosin II, we force the tissue to maximise its global myosin II pool at the apical surface [2] and as a result, lateral cable myosin II declines. These findings highlight how the spatial organisation of the overall myosin II can impact 3D cell shape changes during wound repair. However, the precise *in vivo* distribution and relative contribution of myosin II across apical, lateral, and basal domains remain difficult to resolve experimentally.

To decouple these mechanisms and systematically test their effects, we turned to our 3D vertex model. Based on previous quantifications (Fig. 4, and [2]), we decreased the strength of the purse string by 40% to model shi^TS^, (Supplementary Methods). The model recapitulates the experimental trends in apical wound area closure (Fig. 4B’ and 4O) and apical indentation in both mutants compared to the control (Fig. 4E’ and 4Q). In contrast, to model Mbs RNAi we increased the apical surface tension by an additional 40% and reduced the purse string and lateral cable strength by 21% compared to the in silico control. Whilst the model recapitulated the percentage of apical wound edge cells in Mbs RNAi, the model predicted that shi^TS^ wounds would have less apical wound intercalations than was measured experimentally (Fig. 4D’ and 4P). We hypothesise that tissue-level fluctuations or anisotropies may contribute to the number of intercalations over time. For example, uniaxial cyclic stretching has been linked to T1 transitions [25] and elevated temperatures increased membrane fluctuations and fluidity [26]. Thus, in the absence of such fluctuations or anisotropies in our model, we expect to measure fewer intercalations compared to *ex vivo* conditions such as in the temperature-shifted shi^TS^ experiments.

Together, these findings reveal a critical mechanical balance between purse string tension and background cortical tension. Both impaired purse string assembly via shi^TS^ and excessive myosin II activation via Mbs RNAi disrupt wound closure by stabilising cortical tension, thereby limiting 3D shape change and dynamic myosin II remodelling (Supplementary Movie 5 and Supplementary Movie 6). Our results also establish that lateral cables contribute to intercalation and tissue indentation, with their function compromised when myosin II dynamics are perturbed, as seen in both mutants.

### Basement Membrane Deformation Correlates with Lateral Cable Myosin II Intensity

Building on our findings that lateral cables assemble independently and contribute to apical shape changes, we next explored their role in shaping the basal surface and Collagen-IV-rich BM. To understand how the basal tissue surface deforms during wound healing, we used live imaging of the BM and myosin II at the basal side of wounds over time. We computationally extracted the BM as a single layer, based on Collagen-IV-associated pixel intensities. To provide a tissue scale description of BM deformation, we first quantified the relative change in BM height within the field of view (Fig. 5A). Unexpectedly, on average, we failed to observe any pronounced change in BM height until approximately 60 min after wounding (Fig. 5B), after which the BM gradually deformed, leading to the characteristic domed shape (Fig. 5A) we previously described (Fig. 1C, magenta). We next described local BM deformation by quantifying the mean curvature in 1µmx1µm square bins (Fig. 5C). Similarly to BM height, there was no significant change in total mean curvature during the first hour after wounding. After this point, however, the BM began to bend inward, and its curvature became increasingly negative, particularly around the edge of the wound, indicating that the surface was curving toward the apical side of the tissue (Fig. 5D, 119min). This inward bending coincided spatially with the appearance of lateral actomyosin cables close to the BM surface as revealed by high-power, single-timepoint imaging (Fig. 5E). This suggested that the basal surface, like the apical domain, may be subjected to forces generated by contraction of the lateral cables. To confirm that the pattern of basal deformations indeed matches the spatial distribution of lateral actomyosin cables, we correlated mean curvature with suprabasal myosin II intensity 2 µm above the BM. We found that myosin II intensity was strongly correlated with BM curvature towards the apical surface (Fig. 5F).

**FIGURE 5:**
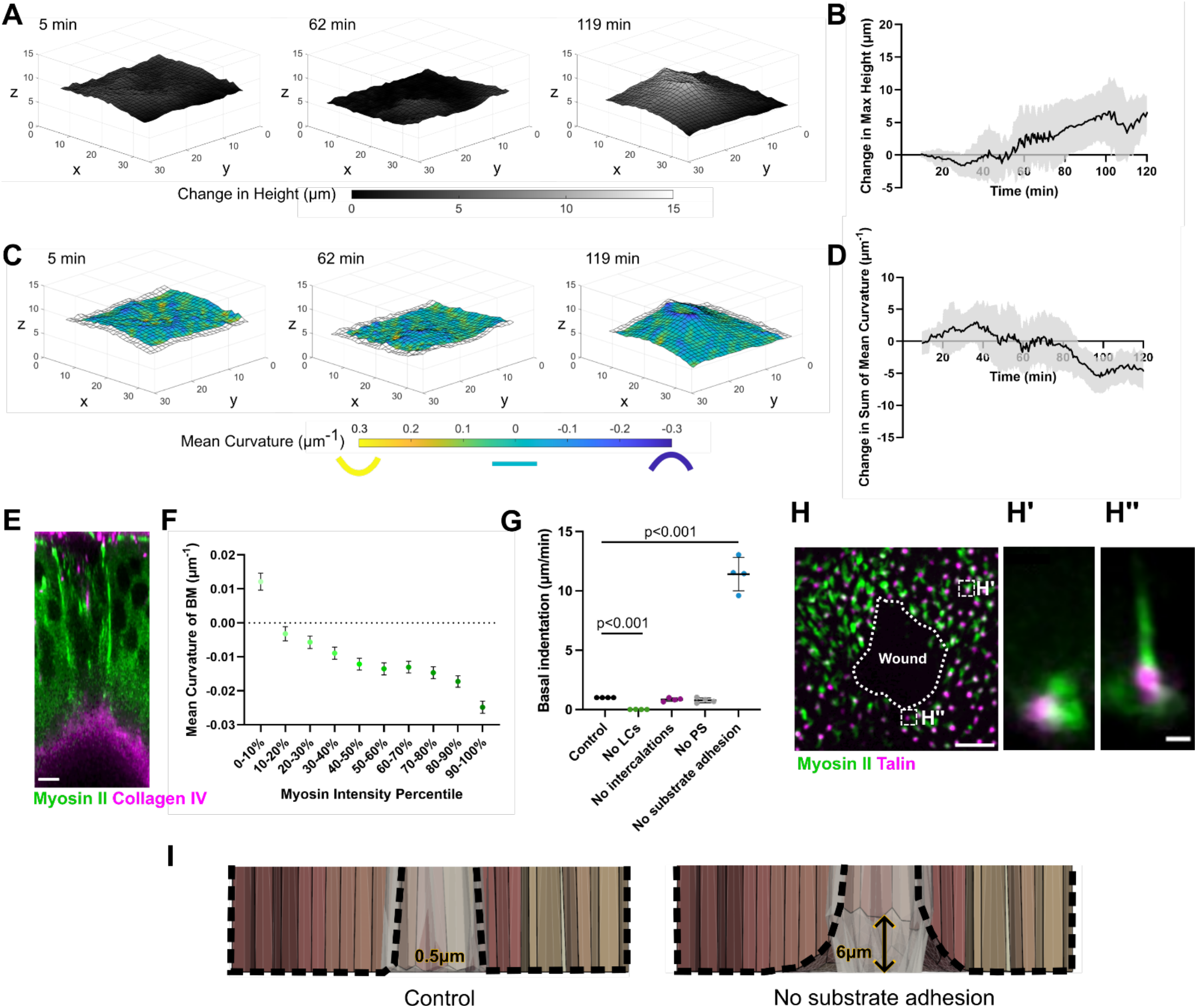
Basement Membrane deformation is delayed and correlates with lateral myosin intensity. A) BM Height over wound closure. B) Change in Max Height. C) Mean BM curvature. Yellow corresponds to regions with positive curvature, light blue corresponds to zero curvature or flat regions, and dark blue corresponds to regions with negative curvature. D) Change in the Sum of Mean Curvature. E) Cross-sectional view of a wounded wing disc expressing sqh-GFP and vkg-mScarlet. Scale bar, 5μm. F) Suprabasal myosin II intensity vs Mean curvature of the BM. G) Basal indentation predictions of the in silico model at 60 minutes. Control (WT) in black, no lateral cables in green, no intercalations in purple, no purse string in grey, and no substrate adhesion in blue. All simulations n=4. H) Max projection of the basal wound surface 60 min after wounding with sqh-GFP (green) and rhea-mCherry (magenta). Scale bar, 5μm. H’) Cross-sectional max projection of a puncta away from the wound edge and H’’) at the wound edge. Scale bar, 1μm. I) Basal indentation predictions of the in silico model at T_end_. Measurements are the average of the control and no substrate adhesion, respectively. G shows individual data points with the mean (horizontal line), SD (bars) and p-values from two-tailed Mann-Whitney tests.

To provide mechanical insight to basal wound dynamics, we used our new 3D vertex model to explore dynamic characteristics that were inaccessible to 2D models, such as the basal tissue indentation or basal wound area during wound repair (Fig. 5G, Fig. S3B-D and Supplementary Movies 1-6). Notably, when a cell intercalates away from the wound edge apically, it may remain at the wound edge basally, giving rise to an increased number of scutoids over time. Interestingly, the 3D vertex model predicted that both apical and basal indentations do not occur without lateral cables, highlighting their essential role in driving cell shortening (Fig. 5G). Furthermore, one of the computational model’s parameters is the strength of substrate adhesion, where we modelled the adhesion as an elastic spring attached to a given Z plane, mimicking an underlying substrate such as the BM (Supplementary Methods). When this adhesion was removed before wounding, basal indentation increased significantly in our computational model as the wound closed (Fig. 5G and Supplementary Movie 7), reaching approximately 6 µm at 60 min (Fig. 5I). This basal indentation is substantially greater than the basal indentation measured in control in silico wounds with a substrate adhesion term reaching approximately 3 µm at 60 min (Supplementary Table 1, Supplementary Movie 1, and 7).

To investigate the adhesion between lateral cables and the basal surface in our *ex vivo* wing discs, we imaged Talin, a key adapter protein that links Integrin adhesion complexes to the actomyosin cytoskeleton at the basal surface [27,28]. We found that myosin II forms a ring around Talin puncta, which pepper the basal surface of the wing disc (Fig. 5H-H’). There were occasional ‘torpedo-like’ myosin II structures projecting away from the basal surface in XZ, which were possible lateral cables (Fig. 5H’’). The colocalization of these myosin II torpedoes with Talin suggests that Talin may serve as anchoring points for the lateral cables.

### Computational Model Predicts Integrin-Adhesion Complex Perturbation Alters Cell Shape

We disrupted the proposed attachment of the lateral cables to the BM by either knocking down Talin using RNA interference (Talin RNAi) or expressing a dominant-negative form of the βPS integrin subunit (Integrin DN) [29]. Because changes in BM height and curvature did not become apparent until 60 minutes after wounding, we extended imaging to 120 minutes post-wounding. Apical wound healing appeared mostly unaffected by Talin knockdown (Fig. 6B-F). However, the mean LC myosin II intensity failed to increase between 10 min and 60 min after wounding, compared to controls, suggesting that Talin may stabilise cable contractility at later time points without being essential for their assembly (Fig. 6G). By 120 min after wounding, the mean LC myosin II intensity decreases to match myosin II levels quantified in the control lateral cables (Fig. 6G). We verified these results using a second Talin RNAi line (Supplementary Fig. 6).

**FIGURE 6:**
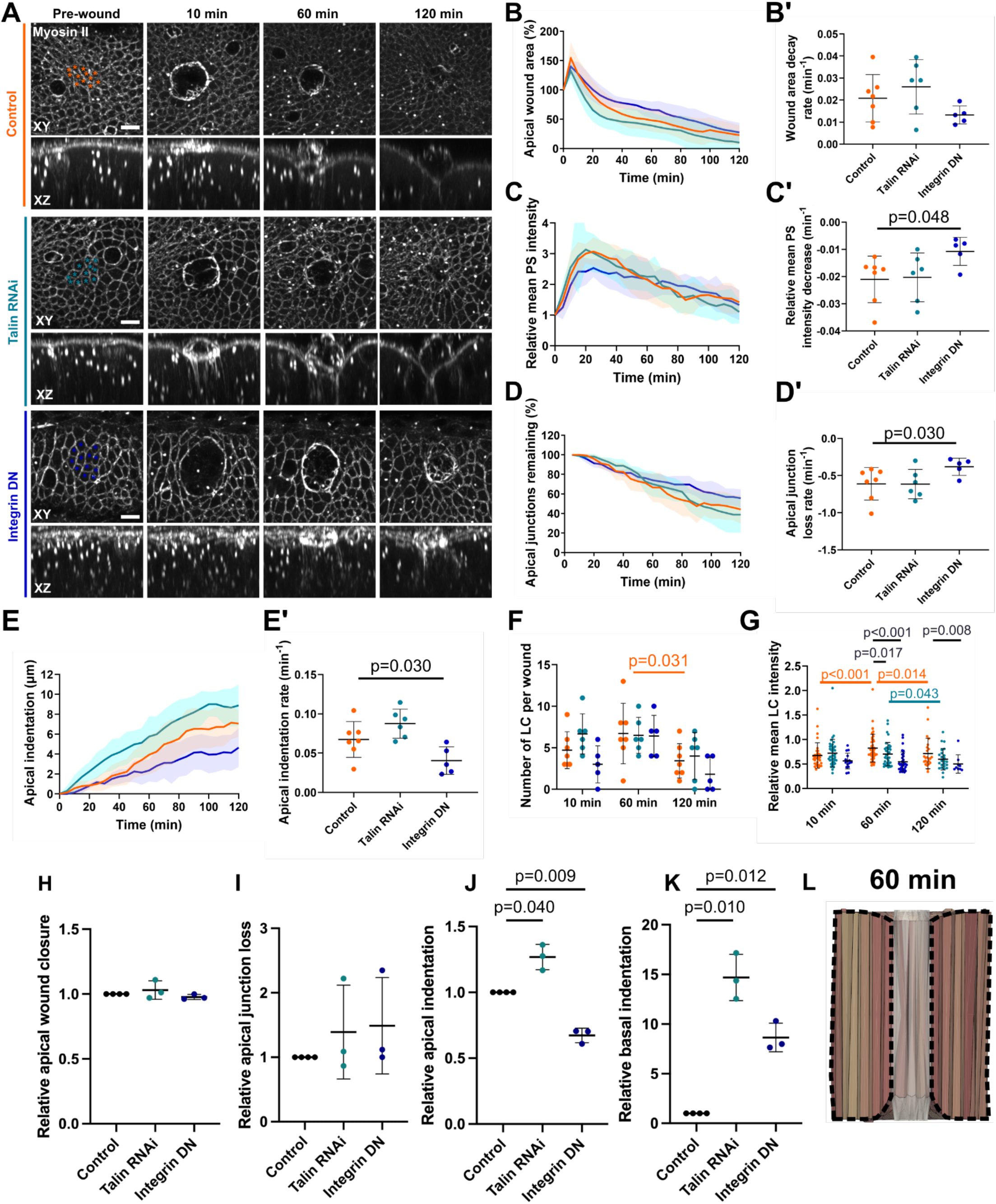
Integrin-adhesion complex perturbation alters myosin II organisation without blocking repair. A) Timelapse images for wounded control discs (orange, n=7), Talin RNAi discs (light blue, n=9) and Integrin DN discs (blue, n=5) expressing myosin II-GFP. Asterisks indicate dead cells. Scale bars, 5μm. B) Apical wound area. B’) Wound area decay rates from fitting one-phase decay curves of individual replicates from B. C) Relative mean PS myosin II intensity. C’) Relative PS myosin II intensity reduction rates from fitting straight lines of individual replicates from C from 20 min after wounding. D) Apical junctions remaining as cells are eliminated from the wound edge. D’) Junction loss rates from fitting straight lines of individual replicates from D. E) Apical indentation. E’) Apical indentation rates from fitting straight lines of individual replicates from E. F) Number of cables per wound 10 min, 60 min and 120 min after wounding. G) Relative mean LC myosin II intensity 10 min, 60 min and 120 min after wounding. H-K) Relative apical wound closure velocity (H), apical percentage of junctions’ loss (I), indentation velocity (J), and basal indentation velocity (K) normalised for each control simulation. Control simulations (n=4, black), Talin RNAi (n=3, light blue), Integrin DN (n=3, blue). L) Graphical representation of the 3D vertex model without substrate adhesion at 60 minutes. Black dotted lines highlight the shape of the tissue. For B-E, shaded areas represent the SD. For B’-E’ and F-G, plots show individual data points with the mean (horizontal line), SD (bars), and p-values from two-tailed Mann-Whitney tests. For F, p-values between different timepoints were calculated using Wilcoxon matched-pairs signed rank tests.

Surprisingly, we found that Integrin DN wounds healed as quickly as controls (Fig. 6B-B’) despite reduced purse string myosin II dynamics (Fig. 6C-C’), apical junction loss rate (Fig. 6D-D’), and apical indentation rate (Fig. 6E-E’). Integrin DN also reduced the mean LC myosin II intensity compared to controls 60 min and 120 min after wounding (Fig. 6G). This suggests that while integrin-based anchorage is not required for cable formation, it may modulate cable function and BM coupling during later phases of repair, similarly to Talin.

As imaging and quantifying BM deformation (Fig. 5) in Integrin DN is experimentally challenging due to the tight folds of the tissues, preventing a clear area surface to extract [30], we instead used our model to test the effect of removing substrate adhesion on *Drosophila* wing disc cells on wound repair. We removed the substrate adhesion and modified lateral cable strength, according to *ex vivo* observations. We assumed all lateral cables apply the same force throughout, and the number of cables is constant over the first hour of closure; we computed the force applied by multiplying the average intensity per cable by the number of cables. Therefore, the force exerted by them in the control would be 3.1 at 60 minutes, Talin RNAi 4.3, and Integrin DN 2.1 (Supplementary Methods). We found that our 3D computational model successfully predicted apical wound area closure in both mutants (Fig. 6H). The model predicted similar apical wound edge cell dynamics between the control and both Talin RNAi and Integrin DN (Fig. 6I), unlike *ex vivo* Integrin DN results. Interestingly, the model predicted that Talin RNAi wounds would apically indent faster than the control, following the same trend as *ex vivo* data (Fig. 6E’ and J). In contrast, the model accurately predicted that Integrin DN wounds would apically indent more slowly than controls. To obtain these predictions, the in silico model required both Talin RNAi and Integrin DN wounds to basally indent more (Fig. 6K-L, Supplementary Movie 8-9) than control simulations (Fig. 3B, Supplementary Movie 1). We hypothesize that basal indentation compensates for the reduced apical indentation rate in Integrin DN wounds to maintain normal wound healing. Due to weakened BM attachment, the basal surface can deform more easily with weaker lateral cables.

## Discussion

Our findings redefine the classical 2D model of epithelial repair by identifying a previously unrecognised contractile system that generates tension in the third dimension. Using the *Drosophila* wing disc, we show that wound edge cells undergo active apicobasal shortening while conserving volume, driving planar expansion into the wound gap. This 3D shape change is driven by the formation of lateral actomyosin cables, contractile structures that span the apicobasal axis and mechanically integrate with the apical purse string.

We confirmed the role of lateral cables using a newly developed 3D vertex model that incorporates substrate adhesion and scutoid geometries as a novelty. Our model simulations demonstrate that coordinated apical and lateral contractility accelerates wound closure, whereas loss of either slows repair. Notably, apical indentation and lateral cable assembly persist even in the absence of an apical purse string, as shown in both the model and experimentally using *shi*^*TS*^. A particularly interesting finding is the prevalence of cell-cell intercalations in *shi*^*TS*^ despite the absence of a purse string. Contrary to previous studies [8,31], these results indicate that intercalations do not depend on apical contractility and cell elongation alone. Our work suggests that changes 3D cell shape, particularly, in the Z axis, such as the lateral cables, may affect the frequency of T1 transitions. The independence of cell intercalation and cell elongation was also suggested by Casani et al., where impaired endocytosis prevented cell-cell intercalation, but not cell elongation [32]. The *in silico* model predicts that loss of lateral cables specifically impairs cell intercalations, emphasizing the importance of the lateral domain in shaping *Drosophila* wing disc during repair. We propose that while the apical purse string primarily drives closure in the XY plane, lateral tension plays a crucial role by facilitating cell volume preservation and aiding cell-cell intercalations. To preserve volume, cells must shorten along the apicobasal axis and elongate along the XY plane. As cells elongate, intercalations help to relax local tissue tension, promoting global tissue remodelling and faster wound closure. This cooperative mechanism offers a conceptual framework for how vertically oriented tissues might efficiently heal in 3D.

Interestingly, although tissue stiffness is known to delay wound closure by restricting apical intercalation [2], weakened BM adhesion under Integrin DN may promote tissue deformation [30]. Normally, wound edge cells are anchored basally to the BM, forcing them to rearrange apically to relieve tissue tension generated by the contracting purse string. When BM adhesion is weakened, as in Integrin DN tissues, basolateral domains gain freedom to deform and accommodate shape changes, compensating for the reduced need for apical rearrangement. We use our model to show that wound edge cells without BM adhesion increase their basal deformation to resume normal closure. These effects may not be needed for wound closure in flatter tissues, such as the previously described *Drosophila* embryo [11]. This freedom to deform likely affects myosin II arrangements, as seen by decreased mean LC myosin II intensity in our Integrin DN wounded tissues. We note that lateral cable myosin II contractility may be inhomogenously exerted along the apicobasal axis. This is challenging to confirm as it is infeasible to accurately quantify myosin II intensity along the apicobasal axis due to the dramatic cell height and its light scattering. The height also presents challenges in performing further nanoablations of the lateral cables to measure recoil in different parts of the cable.

These findings raise further questions about the role of the BM not just as a passive substrate, but as a dynamic mechanical interface that regulates the spatial distribution of tension during tissue repair. Our results suggest that weakened basal adhesion alters cell shape dynamics and mechanical tension distribution, highlighting the BM as an active regulator of 3D wound repair rather than just a passive substrate. We further speculate that BM composition or stiffness could influence the assembly, stability, or function of lateral actomyosin cables and associated remodelling at the basal surface, thereby modulating the transmission of basal tension. However, how extracellular matrix stiffness precisely tunes junctional remodelling remains unclear [33]. To support these insights, we developed a novel, flexible 3D vertex model that integrates scutoid cell geometry and substrate adhesion. This model can be broadly applied to study other epithelial tissues with complex 3D architectures. Altogether, our work expands the paradigm of epithelial wound healing from a 2D phenomenon to a multi-axial, 3D coordinated response.

## Data availability

Data is available upon request.

## Code availability

The code for the 3D vertex model is available at https://github.com/Pablo1990/pyVertexModel/tree/v.1.0.0-Paper.

## Acknowledgements

We would like to thank V. Lachina, and R. Barrientos for their comments. We thank Malik Dawi and Adrià Villacrosa for their contributions on the development of the computational model. S.E.L. was supported by Wellcome Trust grant 225439/Z/22/Z. P.V.-M. was supported by EPSRC grant EP/X03139X/1. Y.M. was supported by the MRC award MR/W027437/1, a Lister Institute Research Prize and EMBO Young Investigator Programme.

## Author contributions

Y.M. and R.J.T. conceived the project. S.E.L. and R.J.T. designed, performed and analysed the ex vivo experiments. P.V.-M. and J.J.M. designed and developed the in silico model. M.Z. performed image analysis. Y.M., R.J.T., S.E.L. and P.V-M. all contributed to writing the manuscript and preparing the figures.

## STAR Methods

### Experimental model and study participant details

Drosophila stocks and crosses were raised on conventional cornmeal media at 25°C or 18°C for shibireTS stocks. Experimental strains are detailed in the key resources table.

### Live imaging

Wing discs were dissected and cultured in dissecting media consisting of Shields and Sang M3 media (Sigma-Aldrich) supplemented with 2% fetal bovine serum (FBS, Sigma-Aldrich), 1% penicillin-streptomycin (pen/strep, ThermoFisher Scientific), 3 ng/mL ecdysone (Sigma-Aldrich) and 2 ng/ml insulin (Sigma-Aldrich). Wing discs were either placed in a glass-bottomed fluorodish apical side down, onto a 0.5 µl line of Cell-Tak (Corning, 354240) (previously dried onto glass-bottom on heat plate set to 29C) with 1mL of dissecting media, or sandwiched between two rectangular coverslips within a dissecting media-filled channel formed by two strips of double-sided tape with 0.1 mm thickness (Tesa 05338) as previously described by Dahmann et al. [4]. Through the latter method, discs were wounded on the apical surface, then flipped over and replaced so that the basal surface faced the objective.

Wing discs were imaged on an inverted Zeiss LSM 880 microscope with a Plan Apochromat 63X oil objective (NA 1.4) or an inverted Leica DIVE with an HC PL APO CS2 63X oil objective (NA 1.4). The LSM 880 Airyscan detector was used in confocal mode for sensitive imaging. For high-resolution snapshots of the basal surface, the detector was used in SR mode with the optimal suggested parameters.

For temperature shift experiments involving shiTS flies and controls, mounted wing discs were incubated on a pre-heated the microscope stage at 31°C for at least 1 hr before imaging and subsequent wounding and imaging were performed at 31°C.

### Laser ablation

Wing disc epithelia were wounded using a pulsed Chameleon Vision II TiSa laser (Coherent) tuned to 760nm with 45-50% laser power. 10-15 cells were ablated per wound with 14 manually specified small circular ROIs. The ROIs were positioned at tricellular junctions in a single z-plane at the level of AJs. The majority of imaging was performed using fluorescently tagged myosin as a marker. AJs were located 0.3-0.5 µm below the strongest myosin signal. For wing discs with cell membranes stained with CellMask or marked by CAAX-GFP, the z-plane is 2-3µm below the strongest apical signal.

Nano ablations of the lateral cables were performed 30 min after wounding. Circular ROIs were drawn over the lateral cables several z-planes below the apical purse string to perform a lateral ablation or in regions of cortex adjacent to the lateral cables to perform a control ablation using a pulsed Chameleon Vision II TiSa laser (Coherent) tuned to 760nm with 45-50% power. Sham ablations were performed at ROIs drawn over the lateral cables with 0% power.

### Quantifications of wound edge cells

#### Image processing and analysis

Image reconstruction and analysis were performed using Fiji/ImageJ [34]. 3D reconstructions were performed in napari [35]. Graphs were produced in GraphPad Prism 10. Figures were mounted in Inkscape 1.1. The following pre-processing steps were performed on all image stacks: rolling ball background, 3D Gaussian blur with a radius of 1.0 and the 3D Correct Drift ImageJ plugin.

#### Clone cell height and volume

A custom ImageJ and Python-based pipeline was used to segment and quantify wound edge clones. This pipeline involved converting the mCherry intensity into a binary mask via Li thresholding, performing repeated dilation and erosion operations to completely enclose any holes and applying the Fill Holes ImageJ plugin. Next, masks were manually corrected in napari [35] to fill any holes that were missed by the plugin and remove any mask regions outside of the clone. Finally, the volume and height of the segmented cells were extracted from the segmented 3D masks.

#### Wound area

The wound edge was manually outlined with the ImageJ Polygon tool, and its area was measured using Analyze/Measure in ImageJ. The area was normalised to the area of ablated cells before wounding, which was denoted as the 0 min timepoint. The wound area over time for each wound was fitted with a one-phase decay (Equation 1) in GraphPad Prism 10 from the timepoint after recoil (3 min or 5 min). The following equation was used:

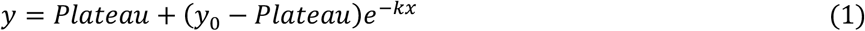

Where x is time, y is wound area starting from y0, and k is the rate constant. The Plateau is constrained to be equal to 0, and k must be greater than 0.

#### Myosin intensity quantifications

To quantify apical purse string myosin intensity, the z-slices containing the purse string, typically 3-4, were maximum projected for each timepoint. Then, the purse string was manually outlined with the ImageJ Polygon tool. The mean intensity was measured using Analyze/Measure. The mean PS intensity over time for each wound was fitted with either a one-phase association (Equation 2) or a straight-line (Equation 3) from the timepoint after the PS intensity peaks in GraphPad Prism 10.

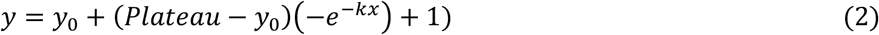

Where x is time, y is mean myosin intensity starting from y0, and k is the rate constant.

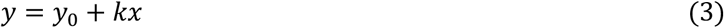

Where x is time, y is mean myosin intensity starting from y0, and k is the gradient of the slope.

To quantify lateral myosin intensity, image stacks were resliced to produce XZ stacks. These XZ stacks were manually inspected for cables, and the corresponding XZ slices for each cable were max projected. On the max projection, a 5 or 10 µm line was drawn over the cable starting from beneath the apical surface signal using the line tool. This measurement also provides the number of lateral cables at each timepoint.

To correct for photobleaching, the relative mean intensities were multiplied by a bleach correction factor. This factor was calculated from the mean apical myosin intensity of unwounded cells at each timepoint normalised by the mean apical myosin intensity before wounding (0 min), referred to as mean resting apical myosin intensity. The mean apical myosin intensity was obtained from a maximum projection of the z-slices containing apical myosin signal within a 100 × 100 px region of interest (ROI), placed at least one cell row away from the wound edge. Myosin intensities were further normalised to the mean resting apical myosin intensity of the border of future ablated cells.

#### Apical junctions remaining

The initial number of wound edge cells was manually counted at the timepoint immediately after wounding. This step was repeated for every subsequent timepoint. The percentage of apical junctions was calculated by dividing the number of wound edge cells at each timepoint by the initial number of wound edge cells. The percentage of apical junctions over time for each wound was fitted with a straight-line (Equation 3, where y is the percentage of apical junctions) in GraphPad Prism 10.

#### Apical indentation

Image stacks were resliced to produce XZ stacks and orthogonal view is used to locate the centre of the wound in ImageJ. Using the Line tool, a line was drawn across the apical surface, and the centroid position was measured. Another line was drawn across the purse string, and the centroid position was measured. The apical indentation is defined as the difference in apical and purse string y centroid positions. The apical indentation over time for each wound was fitted with a straight-line (Equation 3, where y is apical indentation) in GraphPad Prism 10.

#### Statistical analysis

Statistical tests were performed in GraphPad Prism 10. Specific tests and p-values are reported in the corresponding figures.

#### Vertex Model

See Supplementary Methods.

