## Supplementary Methods for "Three-Dimensional Mechanical Cooperativity Optimises Epithelial Wound Healing"

#### 3D vertex model

##### Mechanical equilibrium

We have developed a 3D vertex model that mimics the behaviour of epithelial cells in the *Drosophila* wing disc. The model is based on the minimisation of the total energy of the system ($W$), which is composed of different contributions: volume preservation $W_v$, surface area tension ($W_{sa}$), line tension ($W_{lt}$, used for the purse string and lateral cables mechanisms), substrate adhesion ($W_{sub}$), and an energy barrier that penalises triangular elements with an excessive aspect ratio ($W_{ebAR}$). More specifically, the total energy is a function of the vertex coordinates $\boldsymbol{y} = \{\boldsymbol{y}_1, \ldots, \boldsymbol{y}_{N_y}\}$ and reads

$$W(\boldsymbol{y}) = W_v(\boldsymbol{y}) + W_{sa}(\boldsymbol{y}) + W_{lt}(\boldsymbol{y}) + W_{sub}(\boldsymbol{y}) + \sum_t W^t_{bAR}(\boldsymbol{y}) \tag{1}$$

with the following definitions,

$$W_v(\boldsymbol{y}) = \frac{1}{2}\sum_n \lambda_v \left(\frac{V_n - V_{n0}}{V_{n0}}\right)^2 \tag{2}$$

$$W_{sa}(\boldsymbol{y}) = \frac{1}{2}\sum_n \left(\frac{\sum_f \lambda_{sf} A_f - A_{f^0}}{A^n_0}\right)^2 \tag{3}$$

$$\tag{4}$$

$$W_{lt}(\boldsymbol{y}) = \frac{1}{2}\sum_{\langle i,j\rangle} k_{lt} \left(\frac{l_{ij}}{l_0}\right)^2 \tag{5}$$

$$W_{sub}(\boldsymbol{y}) = \frac{1}{2}\sum_i k_{sub}(y_{zi} - y_{z0})^2 \tag{6}$$

$$W^t_{bAR}(\boldsymbol{y}) = \frac{1}{2}\lambda^t_{AR}\sum_{i=1}^{3}(w^i_t)^2 \tag{7}$$

Here, $n = 1, \ldots, N_c$ is the cell index, $f$ is the face number, and $l_{ij}$ denotes the length of edge connecting vertices $i$ and $j$. $V^n$ is the volume of cell $n$, $V^{n0}$ is the target volume, and $\lambda_v$ is a penalisation factor. In Eq. 4, $A_f$ is the area of face $f$, $A_{f^0}$ is its target area, and $\lambda_{sf}$ is a surface area elasticity parameter.

The term $W_lt$ refers to the line tension energy, where $l_{ij}$ is the length of the edge $(i, j)$, $l_0$ is the target length, and $k_{lt}$ is the line stiffness. In the substrate adhesion energy $W_{sub}$, $y_{zi}$ and $y_{z0}$ denotes respectively the current and target $z$-coordinate of vertex $i$, and $k_{sub}$ is the substrate adhesion.

The energy term $W^t_{bAR}$ measures the aspect ratio of triangle $t$, with $\lambda^t_{AR}$ a penalisation factor. Given the three vertices $\{\boldsymbol{y}_1, \boldsymbol{y}_2, \boldsymbol{y}_3\}$, and denoting by $\boldsymbol{y}_{ij}$ the edge $\boldsymbol{y}_j - \boldsymbol{y}_i$, the aspect ratio is maeasured through the sum of the squares of the three following quantities:

$$\begin{aligned}
w^1_t &= ||\boldsymbol{y_{31}}||^2 - ||\boldsymbol{y_{12}}||^2,\\
w^2_t &= ||\boldsymbol{y_{12}}||^2 - ||\boldsymbol{y_{23}}||^2,\\
w^3_t &= ||\boldsymbol{y_{23}}||^2 - ||\boldsymbol{y_{31}}||^2.
\end{aligned}$$

The equation of motion is found by minimising the regularised total energy,

$$W_\eta = W(\boldsymbol{y}) + \frac{\eta}{2}\sum_i ||\dot{\boldsymbol{y}}_i||^2$$

which is numerically solved by applying and explicit first order time integration scheme,

$$\boldsymbol{y}_{n+1} = \boldsymbol{y}_n + \frac{\Delta t}{\eta}\boldsymbol{g}(\boldsymbol{y}_n). \tag{8}$$

The constant $\eta > 0$ is a friction coefficient, $\Delta t$ is the time step, and $\boldsymbol{g}_i = \frac{\partial W}{\partial \boldsymbol{y}_i}$ is the contribution to the vertex $i$ of the gradient of the total energy $W$.

#### Tissue geometry

##### Initial geometry generation

We construct the initial geometry of the wing disc by creating a 3D mesh of cells with different topology between the apical and basal surfaces. In order to obtain both surfaces, we first segment 2D images from the *Drosophila* wing disc [1] and we use the same image for top and bottom, obtaining, initially, the same topology. Then, several intercalations in the basal side are performed to obtain a given number of apico-basal intercalations (or scutoids [2]). The number of intercalations is equivalent to the number of scutoids in the final geometry. Figure 1a illustrates such construction with 4 cells.

As a result, our description of the cells contains three layers of vertices: top, bottom, and middle [3]. The connectivities of the top and bottom layers represent the topology of the apical and basal surfaces, respectively, while the middle layer is used to connect the top and bottom layers. The latter represents the lateral connections between cells, and in the case of the monolayer at hand, the connections of all cells.

The height of the cells is defined by the parameter of *Cell height ratio* (Table 1), which measures the relative distance between the apical and basal surfaces with respect to the cell diameter.

##### Topology transitions and remodelling

In addition to the vertex coordinates, our geometry description includes a nodal mesh formed by tetrahedra and nodes $\boldsymbol{x}_I$ that are located in the centre of each cell $I = 1, \ldots, N_c$. A segment that joins two nodes $\boldsymbol{x}_A$ and $\boldsymbol{x}_B$ indicates that cells $A$ and $B$ have a common face (see Figure 1b). A similar two-dimensional analogue can be found in [4].

The definition of the external surface of the cells (apical and basal), requires the definition of additional (ghost) nodes that do not belong to any cell. These ghost nodes are connected to each other through a Delaunay triangulation. The particular case of the scutoid is defined by four connected nodes in the middle layer, and thus creating a vertex connecting four cells in the middle of the tissue.

When solving the mechanical equilibrium in equation (8), we include the vertices on the apical and basal surfaces, whose position is computed as the barycentre of the nodes of the tetrahedra they belong to, and additional vertices at the face centres, created when two cells share a face, i.e. there is a segment connecting their center nodes (see Figure 1b). When two tetrahedra share a face, we generate edges connecting the vertices at the centre of each tetrahedra. Note that none of the nodes of the tetrahedral mesh contribute to the total energy of the system, and therefore mechanical equilibrium does not alter the position of nodes $\boldsymbol{x}_I$. This tetrahedral mesh is in fact used to track the topology of the tissue.

| **Parameter** | **Symbol** | **Value** |
|---|---|---|
| Number of cells | $N_c$ | 150 |
| Cell height ratio | | 15/cell diameter |
| Friction coefficient | $\eta$ | 0.07 |
| Basal friction coefficient | $\eta_{basal}$ | $\eta * 600$ |
| Volume penalty | $\lambda_v$ | 1 |
| Target volume | $V_{n0}$ | Initial cell volume |
| Surface area elasticity apical | $\lambda_{sf_a}$ | 1.4 |
| Surface area elasticity lateral | $\lambda_{sf_{lat}}$ | $\lambda_{sf_a} * 0.01$ |
| Surface area elasticity basal | $\lambda_{sf_b}$ | $\lambda_{sf_a} * 0.1$ |
| Target face area | $A_{f0}$ | 0.92$*$ initial face area |
| Target cell area | $A_0$ | 0.92$*$ initial cell area |
| Line tension elasticity | $k_{lt}$ | 0 |
| Target edge length apical | $l_{0_{apical}}$ | 0.0075 |
| Target edge length lateral | $l_{0_{lat}}$ | 0.1578 |
| Target edge length basal | $l_{0_b}$ | 0.0075 |
| Cell shape deformation | $\lambda^e_{AR}$ | 8e-7 |
| Substrate adhesion | $k_{sub}$ | 0.1 |
| Target $z$-coordinate | $y_{z0}$ | -0.24 |
| Remodelling stiffness $R_m$ | | 0.7 |

Table 1: Parameters used for the initial geometry simulation. These values have arbitrary units.

**Remodelling**

At each time step, mechanical equilibrium results in cell deformations, that are eventually followed by topology transitions and remodelling. These transitions are applied according to some geometrical conditions and tolerances. In our case, intercalations are initiated in the apical or basal domain when the relative length of an edge connecting two vertices is shorter than a threshold, which we call *remodelling stiffness* $R_m$.

More specifically, when $l_{ij} < R_m\bar{l}$, with $\bar{l}$ the average edge length of the tissue, we will attempt to establish a connection between the cell-cell nodes or cell-ghost nodes. To do this, we have implemented Algorithm 1, being $A_e$ the set of tetrahedra connected to segment $e$, and $|A_e|$ the cardinality of this set (or segment valence) [6]. Here, *edge* is the connection between vertices, and *segment* connects to nodes from the tetrahedral mesh. Vertices will be moving regarding a gradient $\boldsymbol{g}$, while nodes position would be kept throughout the simulation.

Once we perform the neighbour exchange in the nodal network, we needed to update the geometry of the tissue. Thus, we only updated the vertices associated with the tetrahedra that have changed. Those vertices positions are in the barycentre of the tetrahedra they belong to. Since those vertices might be very different from the previous energy state, we changed them to be as similar as possible to the previous state by moving them a fraction of the distance between the previous and new positions. We then solve the mechanical balance

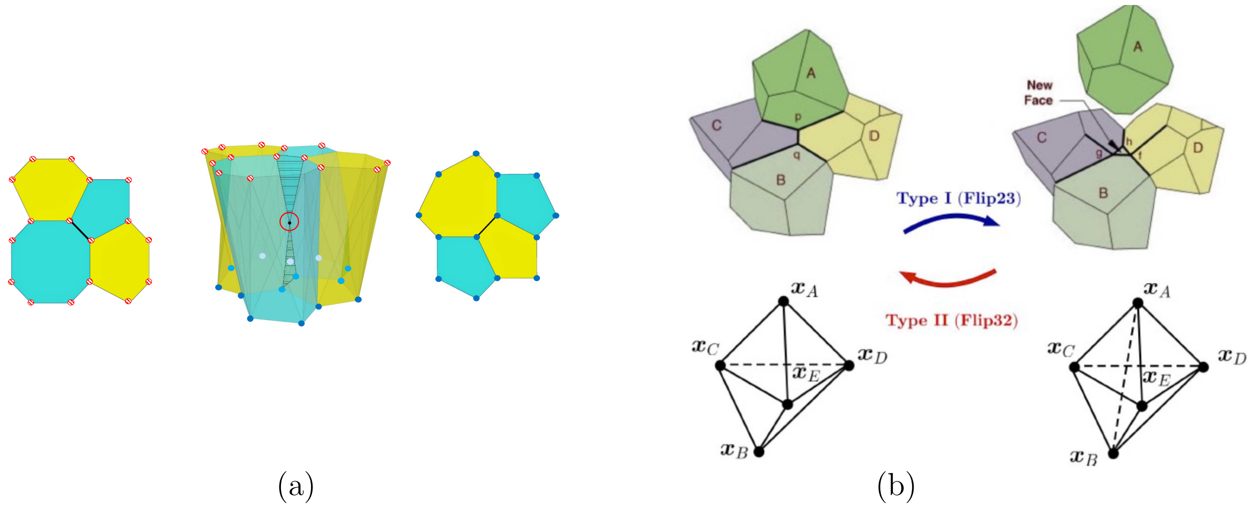


Figure 1: (a) Example of cell geometry construction of 4 cells with topology transition from top to bottom. Left: vertices in top layer (pattern fill). Right: vertices in bottom layer (solid fill). Center: 3D view, with circled vertex belonging to two scutoids. (b) Scheme of the topology transition using a flip 2-3: creation of a new nodal segment (cell-cell connection) from two non-connected cells with centres $\boldsymbol{x}_A$ and $\boldsymbol{x}_B$, resulting in a new segment surrounded by three cells and three tetrahedra. Adapted from [5].

**Algorithm 1** Removal of generic segment

```
if l_ij < R_m l̄ then

    while |A_e| == 3 OR A_e cannot be reduced do
        Perform 2-3 flips to remove faces at segment e
        Update |A_e|
    end while

    if |A_e| == 3 then
        Segment e is removed by a 3-to-2 flip or it cannot be flipped.
    end if
end if
```

of the new configuration (energy minimisation). For that, we run the simulation for a number of time steps to analyse if model will reach an inconsistent configuration ending up breaking the simulation. If it is not, we revert the intercalation.

#### Pre-wound state and boundary conditions

Using the initial geometry, we run the simulation for a number of time steps to reach a steady state, where cells are not changing their shape any more. This is our pre-wound state geometry. In order to mimic a real tissue behaviour, we have used periodic boundary conditions. This means that border cells are connected to their opposite border cells in the $x$ and $y$ directions, simulating an infinite patch on the $xy$-plane.

The parameters used to obtain the pre-wound state geometry are given in Table 1). We note that no fluctuations are introduced in our simulations.

#### Modelling wound healing

To simulate laser ablation, we remove a number of cells from the pre-wound state geometry. In the model, this is done by transforming those *live* cells into a *debris* cell type. In the debris cells, only the energy terms $W_v$ and $W_\eta$ are kept, with a reduced factor $\lambda_v$ indicated in Table 2, while other energy terms are neglected. In addition, purse string ($k_{lt_{ps}}$) and lateral cables tension ($k_{lt_{lc}}$) are modelled via additional line tension $W_{lt}$. For lateral cables in particular, we consider a lateral cable per each edge shared with a debris cell. Note that the resting length for line elements is $l_0 = 0$, while the strength $k_{lt}$ of purse string and lateral cables remains unchanged.

Finally, to model both purse string and lateral cables during wound healing, we follow the evolution from *ex vivo* myosin intensity quantifications, which can be interpreted as a proxy for line tension (main text, fig. 2A).

| **Parameter** | **Symbol** | **Value** |
|---|---|---|
| Number of cells to ablate | $N_{debris}$ | 10 |
| Purse string tension | $k_{lt_{ps}}$ | 3e-5 |
| Lateral cables tension | $k_{lt_{lc}}$ | 7e-5 |
| Debris volume elasticity | $\lambda_{v_{deb}}$ | 1e-8 |

Table 2: Parameters used for laser ablation simulation.

### *In silico* simulations

#### Quantities of interest

We quantified key features of the wound that we use as a descriptor to understand how efficient is wound healing.

- Wound area percentage ($A_w$). We computed the wound area from the total edge length of the contour separating alive cells and debris cells, either at the top or bottom layer. We compare this are with respect to the area of the cells forming the wound before ablation.

- Indentation ($Z_I$). We computed the distance of the cells at the wound edge to a reference Z plane $Z_a = \{(x, y, z)|z = z_a\}$. Different values of $z_a$ where used for the apical and basal side, based on starting configuration. Equivalent units to micrometres.

- Junctions remaining ($N_J$). We calculated the number of cells neighbours of the debris cell.

All these quantities were computed in the apical and basal side.

#### Comparing simulations with different end times

Given that some simulations finished before the 60 minutes, we quantified the velocity of the different measurements regarding that end time $t_{end}$. For instance, we divided the apical indentation by last time step to obtain the "apical indentation velocity". Then, to obtain the relative value we calculate the division between the apical indentation velocity of a given simulation to its correspondent control.

More specifically, if $Q_{sim}$ is the measured quantity in a simulation, and $Q_{sim}^{control}$ the same quantity in a control simulation, we compute the relative rate $q_{sim}$ as,

$$q_{sim1} = \frac{Q_{sim}}{t_{end} * Q_{sim}^{control}} \tag{9}$$

#### Verification of the model

We verified that we can mimic the dynamics of WT wound healing. To better understand how the different mechanisms work, we performed 4 different *in silico* scenarios, where in each simulation we miss one of the following features: purse string, intercalations, lateral cables, or substrate adhesion. This is achieved by modifying the parameters indicated in Table 3. In each simulation, the remaining parameters are kept unchanged.

| Condition | Parameter | Value |
|---|---|---|
| **No lateral cables** | $k_{lc}$ | 0 |
| **No intercalations** | Remodelling stiffness | False |
| **No purse string** | $k_{ps}$ | 0 |
| **No substrate adhesion** | $k_{sub}$ | 0 |

Table 3: Values of the parameters on the different in silico conditions

#### In silico *Drosophila* mutants

In addition, we wanted to understand what mechanics are important in the different *Drosophila* mutants. To do that, we run simulations with the following modified parameters:

| Condition | Parameter | Percentage change |
|---|---|---|
| **Shibire TS** | Purse string tension ($k_{ps}$) | $-40\%$ |
| **Mbs RNAi** | Surface area elasticity apical ($\lambda_{S1}$) | $+40\%$ |
| | Purse string tension ($k_{ps}$) | $-21\%$ |
| | Lateral cables tension ($k_{lc}$) | $-21\%$ |
| **Talin RNAi** | $k_{sub}$ | 0% |
| | Lateral cables tension ($k_{lc}$) | $+38.71\%$ |
| **Integrin DN** | $k_{sub}$ | 0% |
| | Lateral cables tension ($k_{lc}$) | $-32.26\%$ |

Table 4: Parameter changes with percentages for the different mutants.

In particular, Mbs RNAi is calculated from laser ablation measurements related to Figure 5 from [7].

For Talin-RNAi and Integrin DN, we calculated force exerted on average by the lateral cables (main text, Fig. 6) which we calculated by multiplying the number of cables and their associated myosin intensity (3.1 at 60 mins.). We then, did the same calculation for Talin RNAi (4.3) and Integrin DN (2.1). Therefore, the lateral cables strength in Talin RNAi was 4.3/3.1 and 2.1/3.1 for Integrin DN.

### Running the code

The code is freely available at 'pyVertexModel'. For instructions to reproduce the paper code, please use the release 'Paper', where you would find how to create the environment. Once the environment is set and compiled the cython files as explained in the README.md

The steps for running the code are as follows:

1. Obtain the homeostatic geometry by running:

   ```
   python main.py
   ```

   Parameters are set in file parameters/set.py. In this first run, function 'wing_disc_equilibrium' must be changed to the initial *model_name*, being this the file name of the image used as starting topology (e.g. dWL1.tif). Here, there relative number of scutoids in the initial geometry must be chosen (from 0 to 1). The remaining parameters are the ones recommended to obtain wing disc mechanical state.

2. Run 'main.py'. It should run for 80 minutes of simulation time (20 of homeostasis + 60 of wound healing dynamics). After this time, the file 'before_ablation.pkl' will be created, which will be used as starting point for all the conditions mentioned in this article (control, no lateral cables, ...). To obtain those, you have to:

3. Run

   ```
   ./main_paper_simulations.sh
   ```

   Specify the folder in which the simulations are run. This folder should contain the above mentioned 'before_ablation.pkl' file.

For any other question, or issue, it is recommended to create a new issue on GitHub.
